# Compressed sensing for imaging transcriptomics

**DOI:** 10.1101/743039

**Authors:** Brian Cleary, Brooke Simonton, Jon Bezney, Evan Murray, Shahul Alam, Anubhav Sinha, Ehsan Habibi, Jamie Marshall, Eric S. Lander, Fei Chen, Aviv Regev

## Abstract

Tissue and organ function rely on the organization of cells and molecules in specific spatial structures. In order to understand these structures and how they relate to tissue function in health and disease, we would ideally be able to rapidly profile gene expression over large tissue volumes. To this end, in recent years multiple molecular assays have been developed that can image from a dozen to ~100 individual proteins^1–3^ or RNAs^4–10^ in a sample at single-cell resolution, with barcodes to allow multiplexing across genes. These approaches have serious limitations with respect to (i) the number of genes that can be studied; and (ii) imaging time, due to the need for high-resolution to resolve individual signals. Here, we show that both challenges can be overcome by introducing an approach that leverages the biological fact that gene expression is often structured across both cells and tissue organization. We develop Composite *In Situ* Imaging (CISI), that combines this biological insight with algorithmic advances in compressed sensing to achieve greater efficiency. We demonstrate that CISI accurately recovers the spatial abundance of each of 37 individual genes from 11 composite measurements in 12 bisected mouse brain coronal sections covering 180mm^2^ and 476,276 cells without the need for spot-level resolution. CISI achieves the current scale of multiplexing with two orders of magnitude greater efficiency, and can be leveraged in combination with existing methods to multiplex far beyond current scales.

## Main Text

The most commonly used, highly multiplexed methods for imaging RNA are based on singlemolecule fluorescence *in situ* hybridization (smFISH)^11^, with rounds of staining and stripping, and a variety of barcoding strategies to increase multiplexing^4–10^. In “linear” barcoding strategies *(e.g.,* osmFISH^9^ or *in situ* HCR^10^), each color in each round corresponds to one gene. The number of genes *G* that can be measured with *c* colors and *r* rounds is thus *G* = *cr.* In “combinatorial” strategies, such as MERFISH^4,8^ or Seq-FISH^5,6^, gene identity is encoded by a sequence of colors over multiple rounds – that is, many genes may share a given color in a given round, but each gene is encoded by a *unique* sequence of colors across the rounds. The number of genes *G* that can be measured with *c* colors and *r* rounds is thus *G* = 2^*rc*^ −1— ignoring additional rounds needed for error correction. (The number of colors available is typically 2 to 4.)

Multiplex methods have provided an unprecedented tool for tissue biology and histopathology, but they typically measure fewer than 1% of genes, necessitate choosing a gene-expression signature, and can require a week or more to collect these data in a single tissue section. Ideally, it would be possible to quickly generate data on thousands of gene abundance levels in large tissue volumes, perhaps even entire organs.

Notably, existing barcoding methods ignore prior knowledge or biological principles: each spot is decoded independently, without using any ‘local’ information (such as gene expression information at nearby spots). This choice leads to two fundamental limitations on scalability. First, both linear and combinatorial quantification requires imaging at high magnification (up to 100x) so that individual RNA molecules appear as bright, well-separated spots, in order to decode their individual identities from the hybridization images. High-resolution image acquisition over large volumes is a major time bottleneck. Second, there are limitations on the number of genes. In linear barcoding, it is not feasible to substantially increase the number *G* of genes assayed, because the number of rounds of imaging scales with *G* (100-fold more genes requires 100-fold more rounds) and with combinatorial barcoding, increases are limited by optical crowding (spatial overlap between fluorescent spots), because the number of spots scales with *G*. Recent efforts to ameliorate this latter issue with sparser combinatorial barcodes increase the number of rounds of hybridization and, so far, result in a relatively high rate of false positives^12^.

We reasoned that a biology-informed strategy could be more efficient, by incorporating knowledge about the principles of gene expression patterns. Because many genes are co-regulated, measurements of one gene give information about the likely abundances of others. In such cases, one might infer the expression of many individual genes from a much smaller number of composite measurements of gene abundance – mathematically defined as linear combinations of gene abundance levels – consisting of combined signal from multiple genes on the same channel. That is, instead of measuring the level of multiple genes but each of them separately *(i.e.,* each gene in one channel), we should use each channel to measure the composite (sum) abundances of multiple genes in each channel, and later be able to decompress and determine individual gene levels by leveraging the biological insights that genes are co-regulated in modules. We have previously published the theoretical foundations of this strategy, based on the mathematics of compressed sensing^13^, which describes how under-sampled composite data can be decompressed to recover structured, high-dimensional expression signals for individual genes.

Here, we develop such a scheme, Composite *In Situ* Imaging (CISI), implement it in a lab method and computational algorithm, and show that it improves the current throughput of convenient linear barcoding methods by multiple orders of magnitude. Our implementation of CISI consists of four steps (**Fig. 1a**).

**Figure 1.**
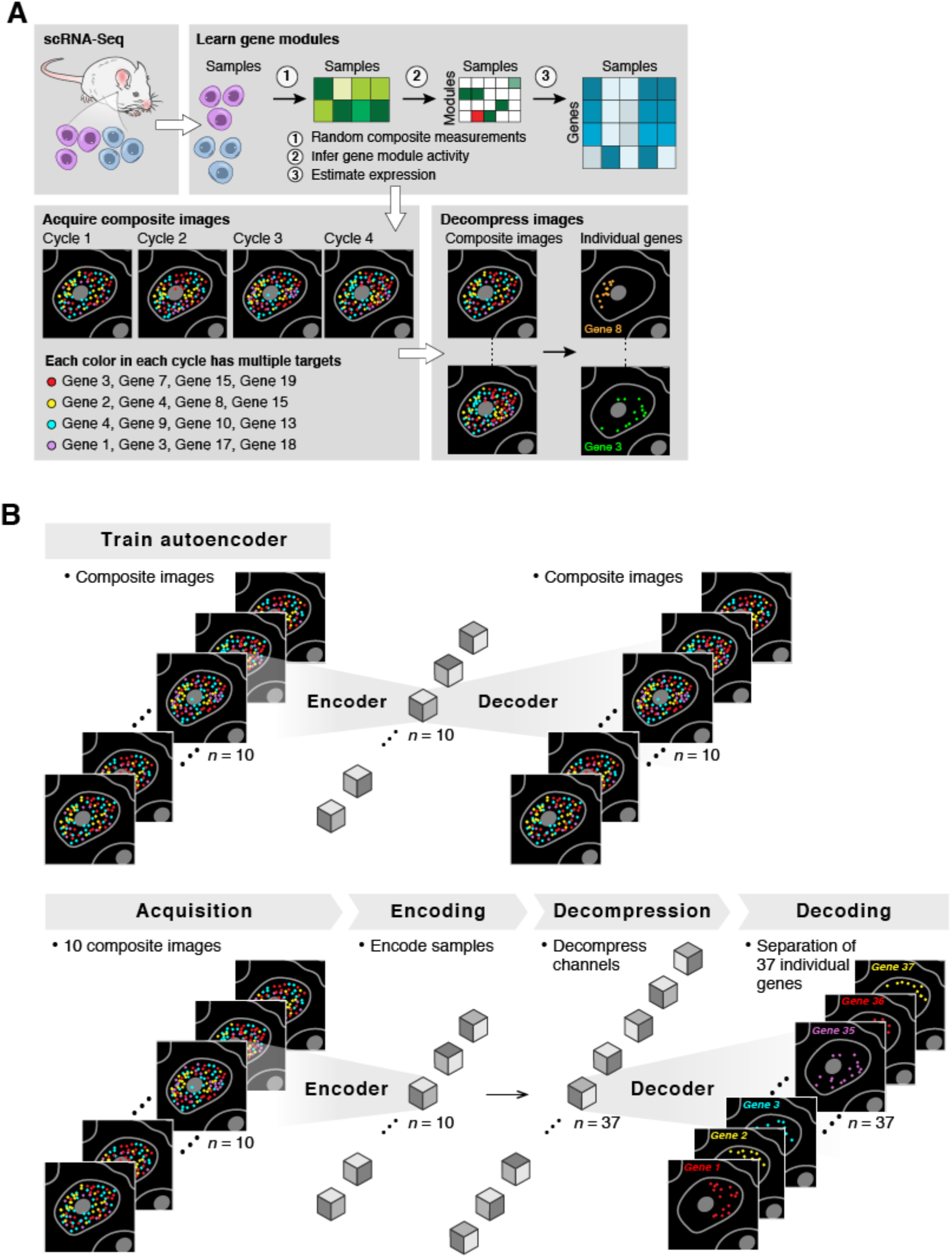
Composite In Situ Imaging (CISI) (**a**) Method overview. snRNA-Seq data (**top left**) is first analyzed (**top right**) to learn a dictionary of gene modules, simulate compressed sensing, and select measurement compositions to be used in CISI experiments (**bottom left**). In a CISI experiment, in each color in each round of imaging, probes for every gene in a given composite measurement are hybridized simultaneously. The process is repeated for different compositions over several cycles of stripping and hybridization (**bottom left**). Finally, composite images are then decompressed computationally (**bottom right**) to recover individual images for each gene. (**b**) Segmentation-free decompression. Top: An autoencoder is first trained on the composite images, with each composite measurement corresponding to one channel. Bottom: Once the autoencoder is trained, the composite images are encoded (“Encoding”), then decompressed to approximate the encoded representation for the unobserved image of each individual gene (“Decompression”), and the pre-trained decoder is used to recover individual images for each gene (“Decoding”).

### (1) Create a dictionary of gene-expression modules

To study the spatial expression pattern of *g* genes in tissue samples, we first obtain single-cell profiles (*e.g.*, from single cell RNA-Seq (scRNA-Seq)) from comparable samples to identify co-expression patterns among the selected genes and to compute a dictionary consisting of *d* sparse gene-expression modules *(i.e.,* each module is a sparse vector of nonnegative coefficients for the genes), such that the single-cell profiles can be well approximated by *k*-sparse linear combinations of the modules (*i.e.*, involving at most *k* non-zero weights)^13^. Below, we use *k*=3 and explain how it was chosen empirically.

### (2) Select composite measurements

We next select *m* composite measurements that enable accurate recovery of the chosen genes. Each composite measurement consists of probes for a subset of genes, corresponding to a linear combination of gene abundance. To select the compositions and numbers of measurements needed for accurate recovery, we simulate compressed sensing in the single-cell data: we first generate a random assignment of genes to measurements, simulate composite measurements as the sum of those genes’ abundances, and compute the recovered (decompressed) profiles. We use simulations to test parameters for (**1**) the total number of measurements *m*, (**2**) the maximum number of measurements in which each gene was included, (**3**) the individual genes for each measurement, and (**4**) the size *d* of the module dictionary and sparsity *k* of the linear combinations. To simplify laboratory implementation, we considered only measurement compositions consisting of binary weights, where each gene was either not included, or included in equal proportion. For each combination of design parameters (**1**) and (**2**), we generate many simulated compositions and compute the recovered profiles from each, and then select compositions that most accurately recover the original expression levels (**Methods**).

### (3) Generate image data

We then synthesize probes for each gene, and create composite probes for each composite measurement by mixing the probes according to the coefficients in the composite design. (Since the weights are binary, the probes for each gene included in a composition are mixed in equal proportions.) We hybridize the composite probes using the linear barcoding approach: in each round, we label *c* composite probes with distinct colors. (For validation, we include one or more additional cycles to directly measure a subset of individual genes.)

### (4) Computational inference of gene-expression in each cell

Finally, we infer the gene expression patterns in the image, using one of two approaches. In the first, the image is segmented into cells. In each segmented cell, we add up the intensity of each color in each round to get a vector y, corresponding to the composite measurements. We then solve a sparse optimization problem to estimate the gene module activities, w, and individual gene abundances, x=Uw, given the composite designs, *A,* and a gene module dictionary, *U.* That is, we solve for w in y=AUw and then calculate x; this is the core optimization problem of compressed sensing. This formulation is more tractable as an optimization problem, because in the modular representation w is sparse, with few non-zero parameters. While the small amount of data we collect, y, is insufficient data to directly estimate all of the parameters of x it is sufficient to infer these few parameters.

In an alternative approach, we analyze the image without cell segmentation or explicit spot detection by using a convolutional autoencoder to infer individual gene abundances at each pixel in the image. Specifically, we use a convolutional autoencoder to compute a low-dimensional, encoded representation of each image, and perform decompression in the encoded latent space (**Fig. 1b, Methods**). In the segmentation-free algorithm we developed for this purpose, we first train a convolutional encoder to represent each of the composite images in a lower-dimensional space. This effectively aggregates local pixel intensities according to data-driven features. At the same time, we train a decoder that can take these encoded representations as input, and then output images that match the originals. Next, for decompression, we take the *m*-channel *encoded* representation of each tissue section as input (each channel corresponding to one of the *m* composite images). For each node in the encoded representation of a given tissue section, we then solve a sparse optimization problem to estimate gene module activities, and compute the encoded representation of the (unobserved) image for each individual gene *(i.e.,* we decompress the encoded representation from *m* to *g* channels). We then decode the encoded representation of each unobserved gene, outputting *g* individual images. During this optimization we include in the loss function the error between the re-composed individual genes and the original composites at both the encoded and decoded layers (among other constraints and regularizations; **Methods**).

CISI offers two important advantages. Like combinatorial barcoding, CISI requires exponentially fewer rounds *r* of hybridization than linear barcoding (r_CISI_ = O(k ln(d)/c), r_combinatorial_ = ln(g)/c, and r_linear_ = g/c; we estimate that, in practice, the number of rounds with either method will be comparable, with r_CISI_/r_combinatorial_ typically between 1/3 and 3; **Methods**). But unlike combinatorial barcoding, CISI does not require spot-level resolution (signal is locally-aggregated – we elaborate on this point in **Discussion** below), and thus allows for faster imaging over large areas: whereas individual spots are often imaged between 60-100x magnification, CISI can be imaged from 10-40x, allowing for imaging that is 2.25-100 fold faster in two dimensional scanning.

To demonstrate CISI, we first piloted it in the mouse primary motor cortex (MOp). We analyzed a set of 31,516 previously published single-nucleus RNA-Seq (snRNA-Seq) profiles from MOp (https://biccn.org/data). We chose to study *g* = 37 genes, consisting of 30 genes that are markers of either broader (excitatory and inhibitory neurons, and various glial cells) or narrower (*e.g.*, layer specific inhibitory neurons) cell subtypes (**Supplementary Table 1**, **Supplementary Fig. 1**), and 7 additional genes that were co-expressed with these markers. In the 27,491 cells in which at least 1 of the 37 genes was detected, the effective number of genes expressed (out of 37, using Shannon Diversity) per cell was 2.87. Given dropouts in snRNA-seq, this is likely an underestimate of true expression.

We then learned a sparse modular representation of the expression of the 37 genes in the 27,491 cells. For cells with only 1 or 2 of the genes expressed, we could trivially represent their expression with 1 or 2 parameters (corresponding to singleton gene modules). For cells with more genes expressed, it is more efficient to represent expression in terms of modules of co-expressed genes. We used our previously published method, Sparse Module Activity Factorization (SMAF^13^), to identify a dictionary of *d* = 80 modules. The modules effectively consist of 1 to 6 genes (2.66 on average; **Supplementary Table 2**), such that the expression of each of the 37 genes in each of 27,491 cells can be represented with a linear combination of 3 or fewer modules with 94.3% correlation. This representation had high correlation with measured expression levels (88%), even in cells with greater than 5 of 37 genes expressed (**Supplementary Fig. 2a**). On average, each cell was described by the activity of 1.72 modules, and most cells were very accurately described (correlation >95%) by just 1 or 2 modules (**Supplementary Fig. 2b**).

We then used the simulation procedure described above to develop barcodes and composite measurements that would allow us to learn the modules in each cell (or in a small region of a tissue section), and subsequently approximate the 37 genes. As expected, performance improved with increasing numbers of measurements (criterion (**1**)), leveling off around 10 measurements (**Supplementary Fig. 3a**). The best compositions using each gene in a maximum of 2, 3, or 4 measurements (criterion (**2**)) resulted in recovered profiles that were 78%, 84%, and 87% correlated with the original profiles (**Supplementary Fig. 3b**). We also considered scalability of probe synthesis. In particular, the number of gene-composition combinations that we would need to synthesize when each gene was included in 2, 3, or 4 measurements was 74, 97, and 120, respectively. Balancing the performance of the different designs and the estimated costs, we selected the best performing set of 10 composite measurements, with each gene included in up to 3 compositions (**Supplementary Table 3**). Using these parameters, we found that simulation refined performance from a median correlation of 76% to 84% with the best performing selection, with these compositions including between 6 and 13 genes (**Supplementary Table 3**).

We synthesized, pooled, and successfully tested the 10 selected compositions. We designed probe pairs targeting multiple regions of each gene for fluorescent *in situ* hybridization with HCR amplification (HCR-FISH^10^, **Methods, Supplementary Table 4**), where each oligonucleotide contains a gene targeting sequence and a barcode that determines the channel (color) of the HCR amplified signal. We assigned each of the 10 compositions to one of three colors, to be imaged during 3 1/3 rounds, pooling the assignment barcoded probes into the 10 compositions. For each of the 10 compositions, we tested these pools by also imaging each gene individually, along with the pool of probes for the composition. We simulated composite images by merging into a composition the images acquired individually for each gene. Taking one composition as an example, the real and simulated composite images agreed well visually (**Fig. 2a**), and had 94.7% correlation between integrated intensity values in segmented cells (**Fig. 2b**). On average, across the 10 compositions the correlation was 90.1%.

**Figure 2.**
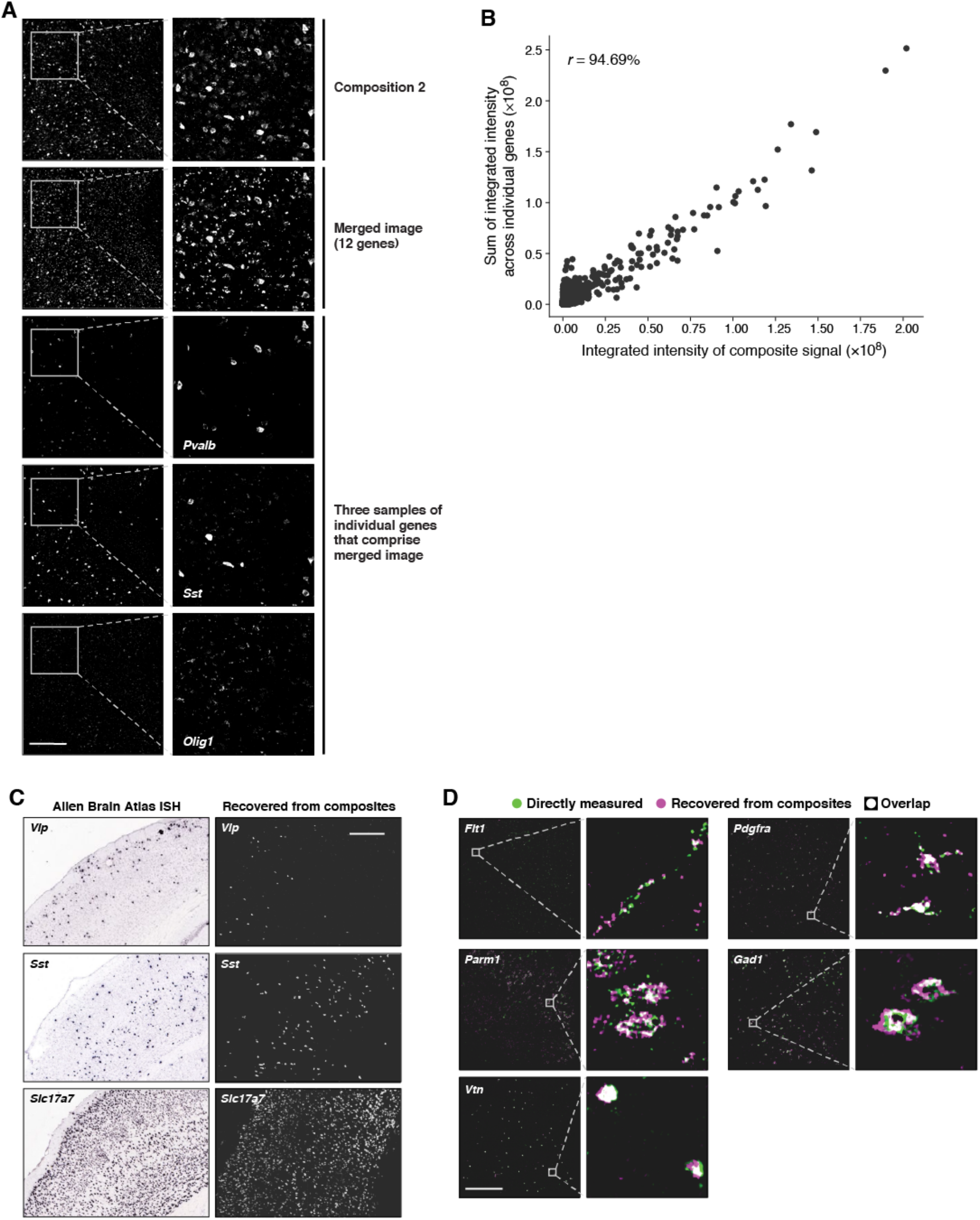
CISI recovers accurate spatial expression patterns from composite experiments. (**a,b**) Quantitative accuracy of the composite imaging. (**a**) A composite image of 12 genes (“Composition 2”) compared to a computational merge of 12 images for each individual gene (“Merged image”). 3 of the 12 individual gene images are shown for reference. Left: entire FOV; Right: zoomed in segment, as indicated. Scale bar: 500um. (**b**) The integrated signal intensity in each segmented cell (individual dots) in the composite image *(x* axis) and the merged image *(y* axis). Pearson’s *r* is noted in upper left corner. (**c,d**) Autoencodeer based decompression successfully recovers accurate spatial patterns of individual genes. (**c**) Agreement with canonical expression patterns. Spatial RNA expression for Vip (top), Sst (middle), Slc17a7 (bottom) by ISH (**left**; Allen Brain Atlas) and in the recovered images by the segmentation free algorithm (**right**). Scale bar: 500um. (**d**) Agreement with individual gene measurements on the same section. RNA images recovered by decompression with the segmentation free algorithm (magenta) and directly measured (green) in the same tissue section. White: images overlap exactly. Scale bar 500um. See also **Supplementary Fig. 4** for additional genes.

Next, we generated a CISI imaging dataset, using our validated composite probe libraries, together with probes for individual genes measured and used only for later confirmation. In each tissue section comprising ~2,500-3,000 cells, we first imaged the 10 composite measurements over 3 1/3 rounds. Using the remaining two colors in the fourth round, and all three colors in a fifth round of imaging, we also directly measured each of up to five individual genes, for subsequent validation purposes. We repeated this in 8 tissue sections, picking different individual genes each time, such that in total we directly measured each of the 37 genes individually along with the compressed measurements (**Supplementary Table 5**).

We decompressed the experimental data with our segmentation-based and segmentation-free algorithms, and evaluated the accuracy of our results in several ways. First, the decompressed images for several genes corresponded well to their known distinct and readily identifiable spatial expression patterns, available as reference images from the Allen Brain Atlas (**Fig. 2c**). For example, these included: Slc17a7, which is broadly expressed in excitatory neurons throughout MOp; Vip, a marker of a subtype of inhibitory neurons, which is expressed more frequently in layer 2/3; and Sst, a marker of another inhibitory neuron subtype, which is expressed more frequently in layer 5 (**Fig. 2c**).

Second, the decompressed images agreed well with the direct measurements of each gene made in the same section, for genes expressed in both rare and common cell types, and in cells of varying morphologies (**Fig. 2d** and **Supplementary Fig. 4**). The correlation between direct and recovered (decompressed) measurements based on integrated signal intensity in segmented cells was high, either when using recovered values from the segmentation-free autoencoding algorithm (83.6%) or when using decompression from segmented cells (88.4%). (We expected the autoencoding algorithm to perform slightly worse by this metric, since it is not optimized for the segmentation masks.) Both are in line with the simulations used to design our measurements, which predicted a correlation of 84%.

Notably, the segmentation-free autoencoder out-performs the segmentation-based algorithm for genes whose expression does not necessarily follow simple patterns (**Supplementary Fig. 5**). The segmentation-based approach omits regions of the image outside of successfully segmented cells, and can result in loss of morphological information, since the output typically consists of filled polygons with uniform intensity. Conversely, the autoencoding algorithm does not omit any regions, and retains morphology by data-driven convolutional features. As a result, while genes like Vtn have expression patterns easily captured by the filled polygons of segmented cells, others such as Flt1 and Parm1 are well-described by autoencoding, but not by segmentation (**Fig. 2d, Supplementary Fig. 5**).

We analyzed cell-type composition of the autoencoding results, by segmenting cells *post hoc* (on decompressed images) and clustering the segmented cells based on the integrated intensity values across genes (**Methods**). Based on the markers in each cluster, neurons comprised about half of all (successfully segmented) cells: 33.3% of cells in 9 excitatory clusters, and 16.9% of cells in 6 inhibitory clusters. In addition, we find 4 clusters of oligodendrocytes and oligodendrocyte precursor cells (16.8%), 3 clusters of astrocytes (12.8%), 3 clusters of microglia (9.1%), 2 clusters of smooth muscle cells (6.5%) and 2 clusters of endothelial cells (4.4%). These *in situ* results are comparable with the representation of these cell subsets in snRNA-seq, albeit somewhat enriched in glial and depleted in endothelial cells: 44.5% excitatory, 14.5% inhibitory, 8.9% oligodendrocyte / OPC, 10.4% astrocyte, 4.6% microglia, 1.9% smooth muscle cells, and 15% endothelial. The under-representation of endothelial cells may be due to challenges in segmenting them (**Supplementary Fig. 5**).

The points of inaccurate recovery were relatively predictable and consistent between the two algorithms, with some false positives for several genes, but few false negatives (**Supplementary Fig. 6a**). Eight genes had some false positive expression patterns in the recovered images that were absent from the direct measurements (**Supplementary Table 6**). In each case, the false positive signals co-occurred with a gene that had probes included in overlapping measurements. For instance, false positives for Hmha1 are found in cells that express Slc17a7 (**Supplementary Fig. 6c, left**). Hmha1 is a member of two compositions, both of which also included Slc17a7, which is additionally included in a third composition.

We developed a simple heuristic to address this, by reducing false positives at the expense of some false negatives. For the 104 pairs of co-measured genes (*i.e.*, that co-occur in more than one composition) that were not correlated (<10%) in snRNA-Seq, we set the expression of one of the two genes to zero whenever they were co-expressed in recovered images (**Methods**). To select which gene to adjust, we calculate the correlation between the 10 composite measurements in a cell, and the pattern of measurements for each of the two genes *(e.g.,* the binary vector indicating which measurements included the gene), and then adjust to zero the gene with the lower correlation. Applying this simple rule reduced false positives and improved the overall correlation from 83.6% to 88.6% with autoencoding, and from 88.4% to 91.6% with segmentation (**Supplementary Fig. 6a,b**).

Using these adjusted values, we found that decompressed measurements are substantially less sparse than snRNA-Seq, while preserving the co-expression programs observed in snRNA-Seq. As previously observed with osm-FISH^9^, the degree of sparsity is much greater in snRNA-Seq than in our decompressed measurements, with 2.87 genes detected on average in each cell in snRNA-Seq *vs*. 5.96 based on decompressed images (6.49 with segmentation) (**Supplementary Fig. 7a**). To compare co-expression patterns, we clustered cells in the *(post hoc* segmented) decompressed data (as discussed above) and in snRNA-Seq (using only the 37 genes), finding 29 and 28 clusters, respectively. Most clusters had expression signatures that were highly correlated with a counterpart in snRNA-Seq, and had identical sets of marker genes *(i.e.,* the gene with the highest normalized expression in each cluster) (**Supplementary Fig. 7b**).

Encouraged by these results, we next generated a substantially larger dataset spanning 12 bisected 2 coronal sections covering 180mm^2^ and 476,276 analyzed cells (**Fig. 3a**). For this experiment we substituted several genes for new genes that would allow us to evaluate performance for both cell type and cell state patterns. These included five immediate early genes (IEGs) that are markers of activity both broadly <u>across</u> cell types (Fos, Egr1) and <u>within</u> specific types (Egr4 in neurons, Klf4 in endothelial cells, Pou3f1 in OPCs)^14^. In addition, we substituted six of our original genes with six cell type markers that are more commonly used by the spatial transcriptomics community (**Supplementary Table 7**; Fgfr3, Tyrobp, Fa2h, Prox1, Sv2c, and Calb1). With this modified panel of 37 genes, we used the Allen Institute Mouse Whole Cortex and Hippocampus SMART-seq (RRID:SCR_019013) dataset to learn gene modules and select measurement compositions. To avoid the earlier co-measured false positives challenges, we increased the number of compositions to 11 and the maximum number of compositions per gene to 4, and expressly selected a design from among top performers in simulation such that there were no exact overlaps, and a minimal number of “large” overlaps (**Methods**). As before, in each tissue section we also individually probed several genes for direct validation (**Supplementary Table 8**). We imaged the tissue at 20X, and decompressed cells using the segmentation-based approach, which, in the current implementation, scales much more effectively than the autoencoding approach. The initial decoded results were less accurate than in our pilot experiment (61.2% correlation with validation images *vs* 88.4% above), which we reasoned was partially due to differences between the training scRNA-Seq and our imaging data – especially expression profiles in the many brain regions that were imaged but not included in training.

**Figure 3.**
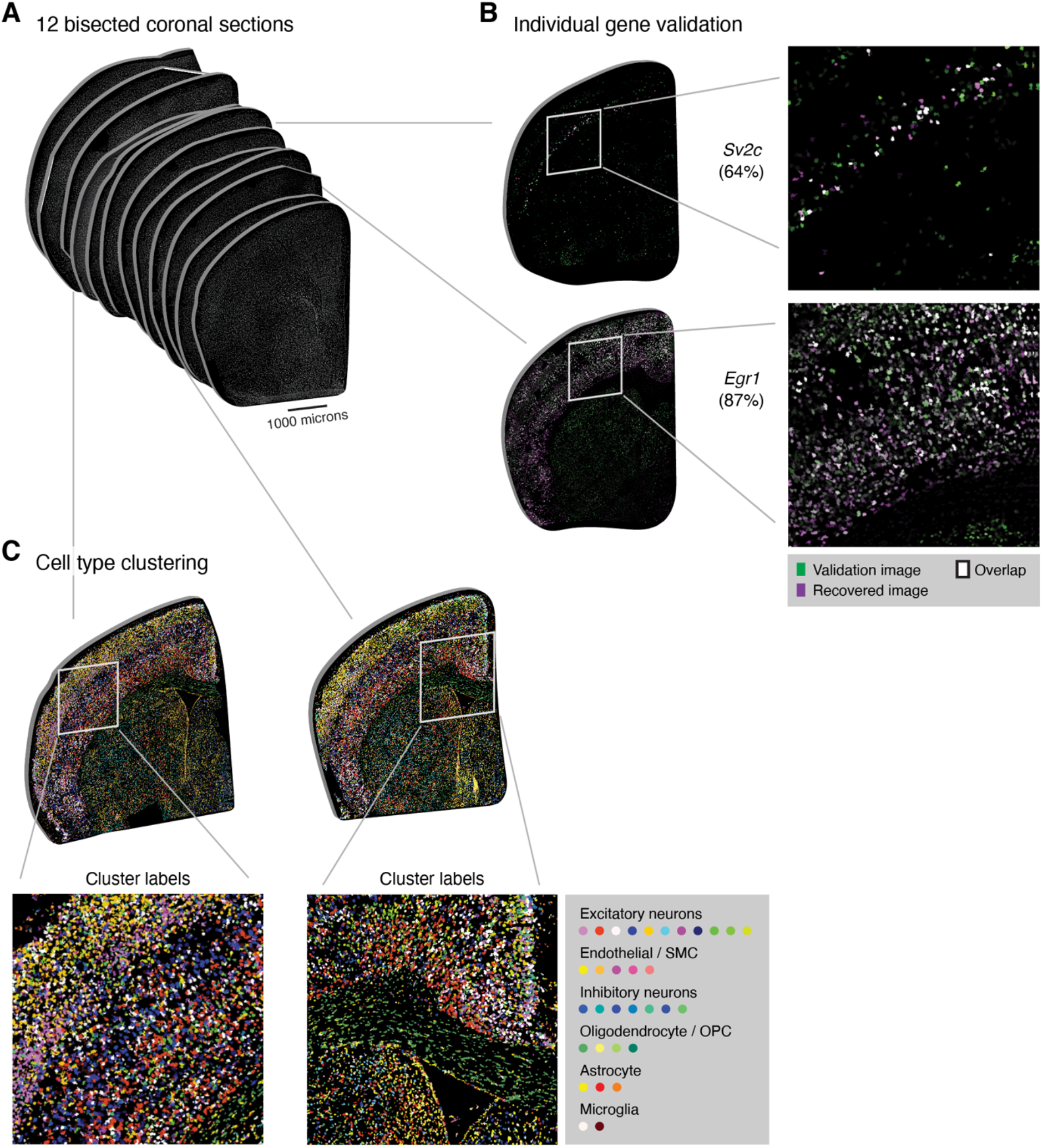
CISI analysis of 12 bisected coronal brain sections efficiently recovers cell type, state, and diverse spatial patterns. (**A**) Composite measurements and validation images were collected in 12 bisected coronal brain sections (greyscale showing DAPI), in total covering 180mm^2^ and 476,276 cells. Composite measurements were decompressed to estimate the expression of 37 genes. Up to four genes were selected for validation in each tissue section. (**B**) RNA images are shown for two genes (Sv2c and Egr1) recovered by decompression with the segmentation-based algorithm (magenta) and directly measured (green) in the same tissue section. White: images overlap exactly. Correlation between recovered and validation images shown in parentheses. (**C**) Cells were clustered into 32 groups (legend) using the decompressed expression levels in all 12 tissue sections. Individual cells are colored based on their cluster label.

We thus developed a computational approach to optimize the module dictionary *post hoc* by using a portion of validation images collected in the new dataset to refine the modules and improve performance. Each cell in the new dataset has validation measurements for up to 4 of 37 genes (with the 4 genes differing in each of 12 sections); we selected 60% of all cells at random in each tissue section, and optimized the modules to minimize reconstruction error in the validated genes in each cell (with the error in ~33 of 37 unvalidated genes in each cell masked out; **Methods**). Using this refined set of modules, we then evaluated performance on each of the validated genes in each section in the remaining 40% of cells, finding it substantially improved with an overall correlation of 81.3% (**Supplementary Table 8**).

The recovered expression levels correctly cluster genes into groups that correspond to known broad and specific cell subtypes (**Supplementary Fig. 8a,b**), with generally good recovery across subtypes (with the exception of Flt1 and Pdgfd in endothelial cells, which suffer from the segmentation issues above). We again find good reconstructions for genes with a broad spatial distribution, such as Slc17a7 or Egr1, and genes with highly-specific spatial expression, such as Sv2c and Tle4 (**Fig. 3b** and **Supplementary Table 8**). On average, the IEGs were recovered at comparable accuracy to other genes, with the worst performance for Pou3f1, the most lowly-expressed IEG (**Supplementary Table 8**). Moreover, the conditional probability of IEG expression is concordant with scRNA and prior findings^14^: a subset of neurons, endothelial cells and OPCs express Egr4, Klf4 and Pou3f1, respectively, while Fos and Egr1 are more broadly distributed (**Supplementary Fig. 8c,d**). This suggests that, with the chosen gene panel, CISI can be used to study cell type as well as cell state. Finally, cell clustering reveals that the recovered spatial patterns correctly capture well known cortical layers and subcortical spatial structures (**Fig. 3c**).

We note several aspects of the results as potential limitations. First, variability in recovery accuracy was associated with relative expression: accuracy was 42% correlated with expression level and 44% correlated with the proportion of cells expressing a gene, such that rarely, lowly expressed genes were less accurately recovered (**Supplementary Table 8**). Second, while existing methods that measure genes individually are more flexible, such that any pattern – or indeed, expression levels with no discernible pattern (spatial or otherwise) whatsoever – can be observed, CISI is constrained to a modular representation of co-expression patterns. However, this representation only applies at the level of individual cells, and thus does not assume any spatial patterning. Furthermore, this doesn’t necessarily mean that CISI can only determine the spatial arrangement of known cell types, because modules can be combined in novel patterns, and the modules themselves are updated using data from the imaged tissue, enabling generalization outside of training data. Nonetheless, it is a limitation relative to other methods. Idiosyncratic expression, unique to one gene, for example due to transcriptional bursting or other inherent variability, may not be well-described by our method.

In conclusion, CISI addresses two key bottlenecks in imaging transcriptomics: increasing the number of genes studied per round of hybridization and decreasing the time needed to scan large tissue volumes per round. Compared with state-of-the-art methods that achieve a similar scale of multiplexing with osmFISH^9^, in the results here we: reduced the number of rounds of imaging 2.5-fold (by assaying 37 genes with 11 compositions plus up to 4 validation images per section); reduced image acquisition time in the xy-plane 25-fold; and reduced acquisition time in the z-plane 8.6-fold. In total, this amounts to a 530-fold gain in efficiency. Much of the gain here is possible because CISI naturally adapts to low-magnification imaging. CISI aggregates signal locally *(e.g.,* within a cell) and decompresses within this local space. If spots within a cell are well-separated their signals are *computationally* summed, at which point the individual spot locations are no longer used. Similarly, when spots overlap their signals are *physically* summed (the pixels do the work of aggregating the intensity for us). In contrast, at low-magnification, optical crowding critically undermines techniques using combinatorial barcodes (such as MERFISH). In principle, CISI could also be used to increase multiplexing with combinatorial labeling (with each combinatorial barcode corresponding to one composite), although high-magnification imaging would be needed to resolve (and decode) each individual spot, and fluorescence crowding would still pose a challenge.

The results here point towards the possibility of greatly increased throughput in imaging transcriptomics. More broadly, CISI is in a class of methods that leverage algorithmic insights and biological structure to be more efficient in generating and interpreting data. Further applications in this class could increase multiplexed protein detection with antibodies, make single cell and single nucleus RNA-seq more efficient by sequencing small pools of cells, or efficiently study genetic perturbations by leveraging common outcomes across experiments.

## Acknowledgements

We thank A. Hupalowska and L. Gaffney for help with figures, S. Farhi, Y. Eldar and members of the Cleary, Chen, Regev, and Lander labs for helpful discussion, and BICCN for open sharing of data pre-publication. Work was supported by the NIH’s *Brain Research through Advancing Innovative Neurotechnologies (BRAIN) Initiative – Cell Census Network* (BICCN; 1RF1MH12128901) (BC, AR and FC) and 1U19MH114821 (AR), Merkin Institute Fellowship at the Broad Institute (BC), Klarman Cell Observatory, HHMI, and NHGRI Center of Excellence in Genome Science (CEGS; RM1HG006193) (AR) and the Eric and Wendy Schmidt Fellows Program at the Broad Institute (FC).

## Author Contributions

BC, FC, and AR conceived the study. BS, JB, BC, EM and AS performed experiments with assistance and feedback from JM. BC, SA, and EH performed snRNA-Seq data analysis, and developed the image processing pipeline. BC developed and implemented the decompression algorithms. BC, AR, FC, ESL, and BS wrote the manuscript with input from all authors.

## Methods

### Mice

All mouse work was done with an adult C57B/L6 mouse according to IACUC procedures specified on protocol 0211-06-18, and we have complied with all relevant ethical regulations.

### Analysis of single-nucleus RNA-Seq data

We selected 37 cell type/layer-specific markers by analyzing snRNA-seq data sets released by BICCN (U19 Huang generated by Regev lab; http://data.nemoarchive.org/biccn/lab/regev/transcriptome/sncell/) for mouse primary motor cortex (M1 or MOp) and generated using the 10x single-cell 3’ protocol (V2). To align the reads, a custom reference was created by 10X Cell Ranger (v.2.0.1, 10X Genomics) using mouse genome and pre-mRNA annotation (Mus_musculus.GRCm38, release 84) according to the instructions provided on the 10X Genomics website (https://support.10xgenomics.com/single-cell-gene-expression/software/release-notes/build#mm10_1.2.0). The default parameters were used to align reads, perform UMI counting, filter high quality nuclei and generate gene by nucleus count matrices. In total, ~30,000 nuclei passed QC metrics including (**i**) the number of unique genes detected in each cell (>200) and (**ii**) the percentage of reads that map to the mitochondrial genome (<10%), and featured in the further downstream analyses using the Seurat package (version 2.2.1).

### Compressed sensing simulations

We use compressed sensing to recover sparse signals from composite measurements. In the basic formulation, we seek to recover sparse gene module activities, *W* ∈ℝ^*d×n*^, and estimate unobserved gene abundances, 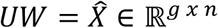 given observations *AUW* = *Y* ∈ ℝ^*m×n*^, a gene module dictionary *U* ∈ ℝ^*g×d*^, and measurement compositions *A* ∈ ℝ^*m×g*^, where there are *m* composite measurements of *g* genes in each of *n* cells, and the dictionary consists of *d* gene modules.

Using the snRNA-Seq data above, and the 37 selected genes, we evaluated different composite designs by simulating composite measurements and recovering individual expression levels by sparse optimization (as previously described^13^; see also https://github.com/cleary-lab/CISI). Briefly, we first randomly selected training, validation and testing subsets, using 60%, 20%, and 20% of all cells for each respective group. In the training set, we calculated a dictionary, *U* ∈ ℝ^*g×d*^, with *d* = 80 modules of *g* = 37 genes (and default SMAF parameters found on https://github.com/cleary-lab/CISI). A given simulation trial with *m* measurements consists of (**i**) randomly assigning genes to compositions, *A*; (**ii**) simulating noisy composite measurements in validation data, *Y* = *A*(*X* + *ϵ*) (with *ϵ* ∈ ℝ^*m×n*^ drawn i.i.d. from a normal distribution with variance adjusted to achieve a signal-to-noise ratio of 5); (**iii**) decoding sparse module activity levels, 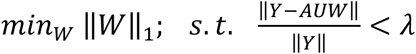; (**iv**) estimating individual expression levels, 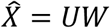; and (**v**) calculating the correlation between the original and estimated levels, 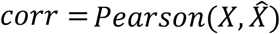. When evaluating different measurement designs in step (**i**), we varied the total number of measurements (from 8 to 12), and the maximum number of measurements in which each gene appeared (either 2, 3, or 4). Each gene was then randomly assigned to a randomly chosen number of measurements (up to the maximum). Final assignments resulting in either two or more genes being perfectly co-assigned or in large measurement imbalance (any gene appearing more than 4 times more frequently than any other gene) were excluded. We then iterated steps (**i**)-(**v**) 2,000 times, selected the 50 composition matrices resulting in the top correlations, and evaluated (steps (**ii**)-(**v**)) in testing data. The correlations in testing data were then used to compare different numbers of measurements, and maximum assignments per gene (**fig. S3**).

### Selection of the final library

For the initial study, based on the comparisons in **fig. S3**, and considering the number of probes that would need to be synthesized in each scenario, we selected the composition with the highest performance in testing data among those with 10 measurements and a maximum of 3 assignments per gene. Each of the 10 compositions was assigned to one of three colors, to be imaged during 3 1/3 rounds (**table S2**).

For the larger study with 12 coronal sections we simulated compositions with 11 genes and 4 assignments per gene to reduce the number of exact overlaps *(e.g.,* where one gene is assigned to 2 compositions, and those 2 compositions match 2 of 3 assignments for a second gene). From among the top 50 performing compositions in simulation with no exact overlaps, we selected the composition which had the fewest number of large overlaps, where a gene has a large overlap if more than 66.7% of its assignments matched the assignments for at least one other gene.

### Probe design and validation

For each target mRNA, HCRv3.0 DNA probe sets of ~20 probe pairs each were ordered from Molecular Technologies, or designed in-house and ordered from IDT. All HCR v3.0 reagents are available from Molecular Instruments, Inc. (molecularinstruments.com). Target binding site sequences can be found in **table S4**.

### Tissue preparation and brain extraction

An adult C57B/L6 mouse was perfused with ice-cold PBS (10010023, ThermoFisher Scientific) prior to dissection of the brain. The brain was then extracted and flash frozen in liquid nitrogen. After OCT embedding, the brain was sectioned directly into an APTES coated 24-well glass bottom plate (82050-898, VWR). For coating, plates were coated with a 1:50 solution of APTES (440140, Sigma) in 100% Ethanol (V1016, DeconLabs) for 5 minutes followed by 3x washes with 100% ethanol before drying. Tissues were fixed in 10% Formalin (100503-120, VWR) for 15 minutes and washed with PBS before overnight permeabilization with 70% ethanol. Tissues were re-hydrated with PBS prior to hybridization. In the larger study, we added a tissue clearing step: re-hydrated sections were cleared briefly with two five-minute washes of 4% SDS (BM-230, Boston BioProducts) prior to hybridization.

### *In situ* hybridization

*In situ* HCR version 3.0 with split-initiator probe sets was performed using the protocol detailed in^10^ with some slight adaptations. Probe sets for each individual target mRNA were diluted to the concentration specified in the protocol and organized into composite channels. A composite channel is comprised of a mix of probe sets for approximately 10 different target mRNAs, each with the same initiator. In total, 10 composite channels were created. Three composite channels can be hybridized per round of imaging. Thus, for the first round of hybridization, probe sets for three composite channels with distinct initiator sequences were added at once to each tissue.

Probes were hybridized for approximately 8 hours in hybridization buffer and then tissues were washed 3 times with 30% probe wash buffer for 10 mins each at 37C and 1 wash of 5X SSCT for 10 mins at room temperature (buffer compositions available from Molecular Instruments). Snap-cooled hairpins were added at a 1:200 diluted concentration and amplification was allowed to proceed for 8 hours. Excess hairpins were then washed off with 3 washes of 5X SSCT (15557044, ThermoFisher Scientific), for 5 minutes each. Tissues were stained with DAPI (1:5,000 TCA2412-5MG, VWR) immediately prior to imaging. After imaging, probes were stripped from tissues using 80% formamide at 37°C for 60 minutes. This entire process (hybridization, amplification, imaging, stripping) was repeated for up to five rounds of imaging (see **table S5** for composites and individual targets imaged in each round). All DNA HCR amplifiers (hairpins), hybridization buffers, wash buffers, and amplification buffers were ordered from Molecular Technologies. All HCR v3.0 reagents are now only available from Molecular Instruments, Inc. (molecularinstruments.com).

### Imaging

Imaging was performed on a spinning disk confocal microscope (Yokogawa W1 on Nikon Eclipse Ti) equipped with a Nikon CFI APO LWD 40x/1.15 water immersion objective or a 20x 0.8 plan apo lambda objective operating NIS-elements AR software with Andor Zyla 4.2 sCMOS detector. DAPI fluorophores were excited with a 405nm laser, Alexa 488 HCR amplifiers were excited with a 488nm laser with 525/36 emission filter (Semrock, 77074803), Alexa 546 HCR amplifiers were excited with a 561nm laser with a 582/15 emission filter (Semrock, FF01-582/15-25), and Alexa 647 HCR amplifiers were excited with a 640nm laser with a 705/72 emission filter (Semrock, 77074329).

### Image processing

Before downstream analysis, we ran a series of image processing steps to normalize, stitch, align, and segment the images in each color, field of view, round, and tissue. In the pilot study, we first took a maximum projection across the z-axis, and then used the DAPI channel to stitch the fields of view within each round of imaging (using ImageJ software^15^). We applied the stitching coordinates from the DAPI channel to each of the other channels. We then smoothed the image for each channel using a median filter (with a width of 8 pixels). (If spot-level resolution is needed, this step may not be advised. Since we do not need this resolution, we use this step to make autoencoder reconstruction an easier task.) From each smoothed image, we aligned and subtracted background signal, obtained by imaging after stripping the final round of fluorescent probes. We then adjusted brightness and contrast by rescaling according to upper and lower thresholds determined using auto-adjust in ImageJ. The same rescaling parameters for each channel (determined from the maximum upper threshold and minimum lower threshold) were applied to all tissues and rounds. After rescaling, we applied a flat field correction to each field of view, by normalizing (dividing) each pixel by the median smoothed pixel intensity across all images (with smoothing by a Gaussian filter with a width 1/8 of the image dimension). Each round of the flat field-corrected images in a given tissue was then aligned using ImageJ. These images were used in the remainder of downstream analysis.

In the larger dataset we used a modified procedure (detailed here: https://github.com/clearylab/CISI). Briefly, raw images for each section and round were first stitched using the BigStitcher plugin for ImageJ^16^. The Starfish pipeline^17^ was used to remove background and call spots (allowing for the detection of large, closely-spaced “spots” to account for low-magnification imaging) in each stitched image. The output was used to create masks and produce filtered images with background removed. Filtered images were aligned across rounds within each section. In some sections, either the stitching or alignment failed in a portion of the section in one or more rounds. These regions were masked and excluded from downstream analysis by comparing stitched and aligned DAPI images from each round, and manually drawing masks around the affected regions.

For segmentation, we used CellProfiler^18^, and calculated one image mask per nucleus in each tissue using DAPI in the first round. Each mask was then expanded by up to 10 pixels (without overlapping a neighboring cell). Comparisons and decompression with segmented cells were done using the integrated image intensity in each expanded nucleus mask.

### Decompression of composite signals

We developed two methods to decompress composite signal intensities into signals for individual genes.

The first method, which we used primarily as a point of reference for validation statistics, is based on cell segmentation. Given the intensities of each composite measurement in each segmented cell, *Y* ∈ ℝ^*m*×*n*^, we solved a sparse optimization problem to decode sparse module activity levels, 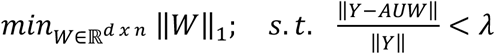, before estimating individual expression levels, 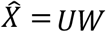, with the same method as in our simulations (above).

The second method we developed decompresses entire images using a convolutional autoencoder. In this approach, for a given set of 10-channel composite images, we first train a model to identify a reduced (encoded) representation of each image, which can then be decoded to recapitulate the original. During this training, we optimize the following loss function:

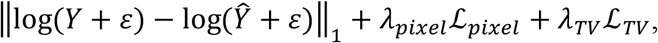

where *Y* is the original image, 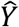 is the decoded image, 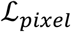 is a loss on pixel density, 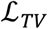 is the total variation of 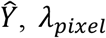 and *λ_TV_* are hyperparameters, and *ε* is a small constant. 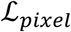 is calculated as the Poisson log-likelihood of the pixel density, which is computed as the Shannon Diversity across pixels, divided by the number of pixels, with prior density set by a parameter *δ_pixels_*. Convolutions in each layer of the network are computed across filters (or kernels), but not across the 10 composite channels. Hence, each of the 10 channels remains separated from the other channels throughout each layer of the network. However, only one set of convolutional weights is learned; these are shared across all channels. The number of parameters in the model is, thus, relatively small, and the autoencoder trained quickly on our data. As discussed below, hyperparameters, including the number of encoding and decoding layers, the number and size of filters, and pooling sizes are chosen by hyperparameter tuning on a small set of validation images.

Using the trained autoencoder, we decompress composite images as follows. First, we encode each 10-channel image to a reduced representation, 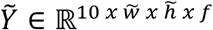, where 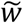 and 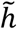 are the reduced width and height (after pooling at each encoding layer), and *f* is the number of convolutional filters. We then solve for sparse module activities, 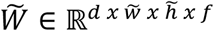, where *d* is the number of modules in the dictionary (here, 80), and then estimate the encoded representation of each individual (unobserved) gene, 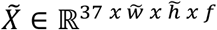. These representations are then run through the pre-trained decoder to produce an image for each gene, 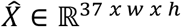.

Our loss function has components at both the encoding and decoding layers:

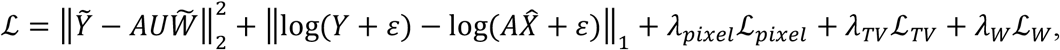

where 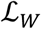 is a loss on the density of 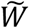, calculated as the Poisson log-likelihood of the Shannon diversity, with prior density set by parameter *δ_w_*. We implemented our model in tensorflow, using the Adam optimizer.

Hyperparameters and model architectures were chosen by hyperparameter tuning on a small set of validation images. The validation images consist of 4 of 36 patches from each of 3 images *(i.e.,* from 3 tissue sections, each with 36 patches), for a total of 12 patches (equivalent in size to 1/3 of one image). Each patch includes signal from the 10 composite measurements, along with up to 5 directly measured genes. In each validation trial, we select hyperparameters, train the autoencoder on the composite data, decompress all genes, and then calculate the trial score as the correlation between the subset of directly measured and recovered genes (this is done in *post hoc* segmented cells, when using the autoencoder). We selected the hyperparameters from the best performing trial, and used these to run our analysis on the full dataset. More details can be found at https://github.com/cleary-lab/CISI.

We applied a heuristic correction to co-measured genes. We first identified 104 pairs of co-measured genes (*i.e.*, that co-occur in more than one composition) that were not correlated (<10%) in snRNA-Seq. For each pair, we set the expression of one of the two genes to zero whenever they were co-expressed in recovered images. To select which gene to adjust, we calculate the correlation between the 10 composite measurements in a cell, and the pattern of measurements for each of the two genes *(e.g.,* the binary vector indicating which measurements included the gene), and then adjust to zero the gene with the lower correlation.

### Optimization of the module dictionary using validation images

For the segmentation-based approach, we developed a method to update the module dictionary, U, using the validation images collected in each tissue section. We initialized the dictionary with the dictionary found from scRNA-Seq training data. In each iteration of this method, we first use the current dictionary to decode sparse module activity levels, W, as described above (Decompression of composite signals). We then update U via gradient descent (as implemented by the AdamOptimizer function in TensorFlow) on the following loss function:

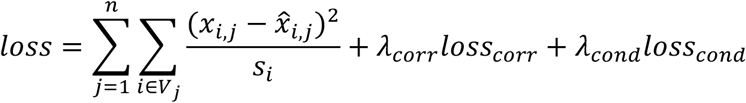

where *V_j_* is the set of up to 4 genes validated in each section and *s_i_* is the scale of each gene (defined as the 75^th^ percentile of nonzero expression). The first term penalizes reconstruction error, while the second and third penalize differences from scRNA-Seq training data in correlation structure and conditional probabilities, respectively. Specifically,

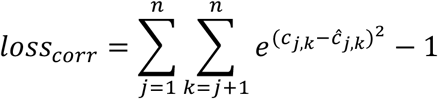

and

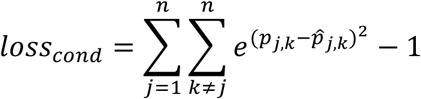

where *c_j,k_* is the correlation between genes *j* and *k* in scRNA-Seq, *ĉ_j,k_* is the correlation between genes *j* and *k* in decompressed results, and *p_j,k_* and 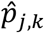 are the probabilities of gene *k* expression conditioned on gene *j* in scRNA-Seq and CISI respectively. The exponential form of these penalties was chosen so that the loss would be similar to the squared deviation for small differences, while scaling much faster for large differences. The method was run for 25 epochs with 5,000 cells per iteration batch.

When updating the dictionary in this way we first partitioned the cells into training (60%) and testing (40%). To find optimal hyperparameters (*Λ_corr_* and *Λ_cond_*) we perform a grid search over 100 pairs of parameters. For each pair, we update the dictionary using training cells, and then use this dictionary to decompress composite measurements in test cells. We select the pair of hyperparameters for which performance (here, average gene-wise reconstruction correlation) rapidly decays when increasing either parameter individually (*Λ_corr_* = 1.0 and *λ_cond_* = 2.15).

### Plotting decompressed images

The decompressed results for each gene vary in their relative signal intensities (as do direct measurements for each gene). When plotting merged validation images (as in fig. 2d and fig. S4), we normalize the signal for each gene to automatically adjust contrast and brightness. The specific parameters of this normalization can be found in the code demo of our online repository (https://github.com/cleary-lab/CISI/blob/master/getting_started/plot_decompressed_images.py). The signal plotted for direct measurements have been pre-processed according to the methods described above.

### Comparison of CISI and combinatorial barcoding

We can approximate the number of imaging rounds in CISI, rCISI, relative to that in combinatorial barcoding methods, r_combinatorial_, defined as r_CISI_/r_combinatorial_, based on our results here, using simulation to extrapolate to larger scales, and by comparing with existing combinatorial methods. Here, we used 3 and 2/3 rounds to measure 37 genes; the same could be achieved using 3-color combinatorial barcoding without error correction. More commonly, 4 or 5 rounds would be used to measure 37 genes and allow for error correction. At larger scales, simulations in our earlier work^13^ suggest that ~100 composite measurements would suffice to approximate the expression of 10,000 genes. This could be done in 33 and a 1/3 rounds of CISI. To date, the only combinatorial method to scale to this level did so with 80 rounds of imaging^12^. We therefore *very roughly* approximate that the required rounds of imaging with either approach will be comparable, and that r_CISI_/r_combinatorial_ will be in the range 1/3 to 3, allowing for improvements in combinatorial methods and the possibility of needing more rounds than anticipated with CISI.

## Data availability

We used publicly-available snRNA-Seq data sets released by BICCN (U19 Huang generated by Regev lab; http://data.nemoarchive.org/biccn/lab/regev/transcriptome/sncell/), and full-length scRNA-Seq (the Allen Institute Mouse Whole Cortex and Hippocampus SMART-seq (RRID:SCR_019013)).

## Supplementary Figure legends

**Supplementary Figure 1.**
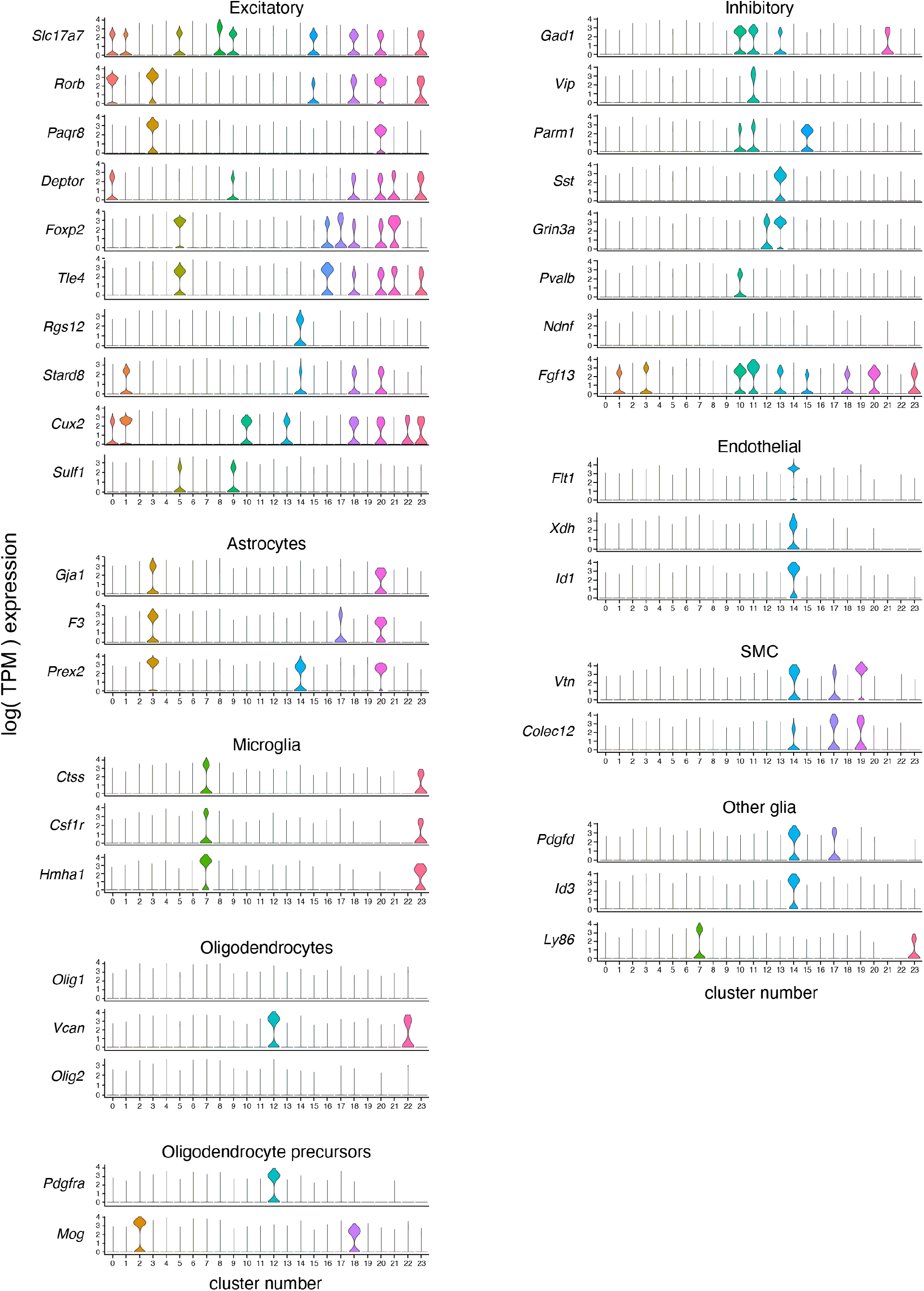
Marker gene expression in snRNA-seq clusters. For each of 37 genes, shown is the distribution of expression (individual violin plots; y-axis) in each of 23 snRNA-Seq clusters (*x* axis). Marker genes for similar cell types are grouped together with the cell type labeled on top.

**Supplementary Figure 2.**
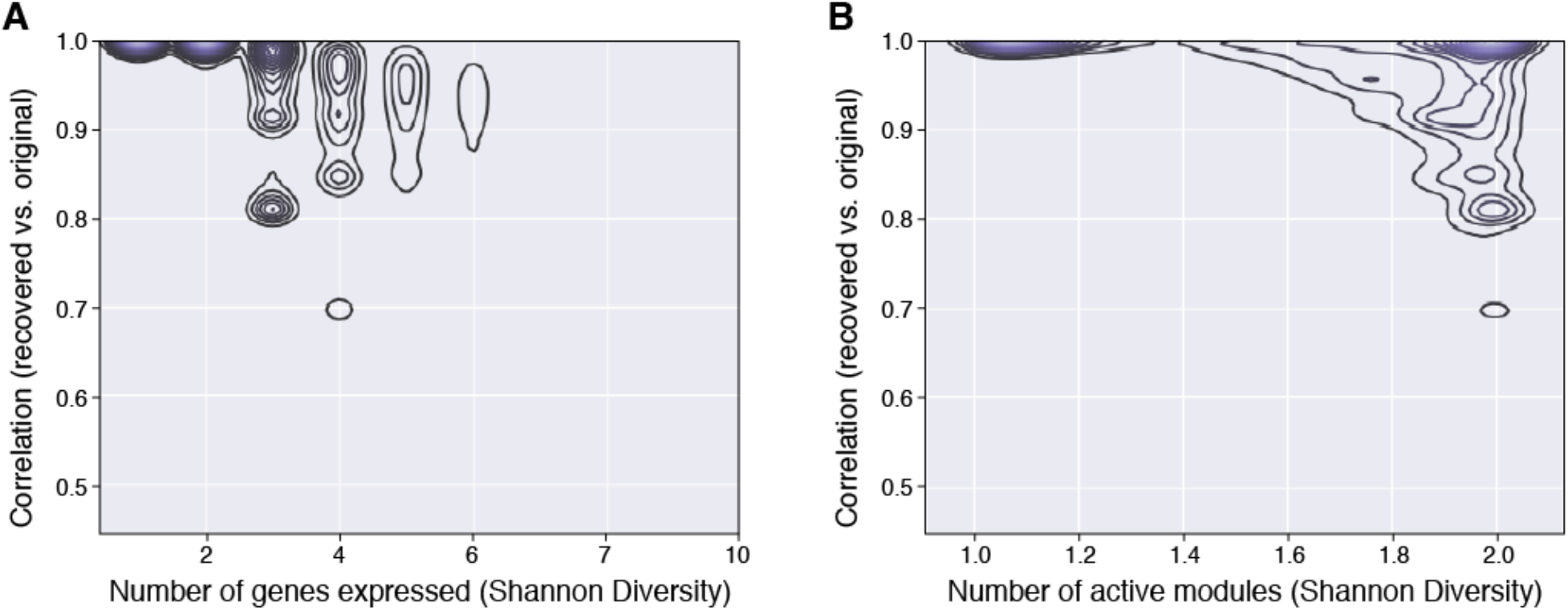
Analysis of modular factorization based on gene and module diversity. Pearson correlation (y-axis) between the original expression levels of 37 genes in each cell and those approximated in those cells by Sparse Module Activity Factorization (SMAF). Contour plots depict the density of cells at each level of correlation with either a given number of genes expressed (**a**; x-axis) or a given number of gene modules by SMAF decomposition (**b**; x-axis).

**Supplementary Figure 3.**
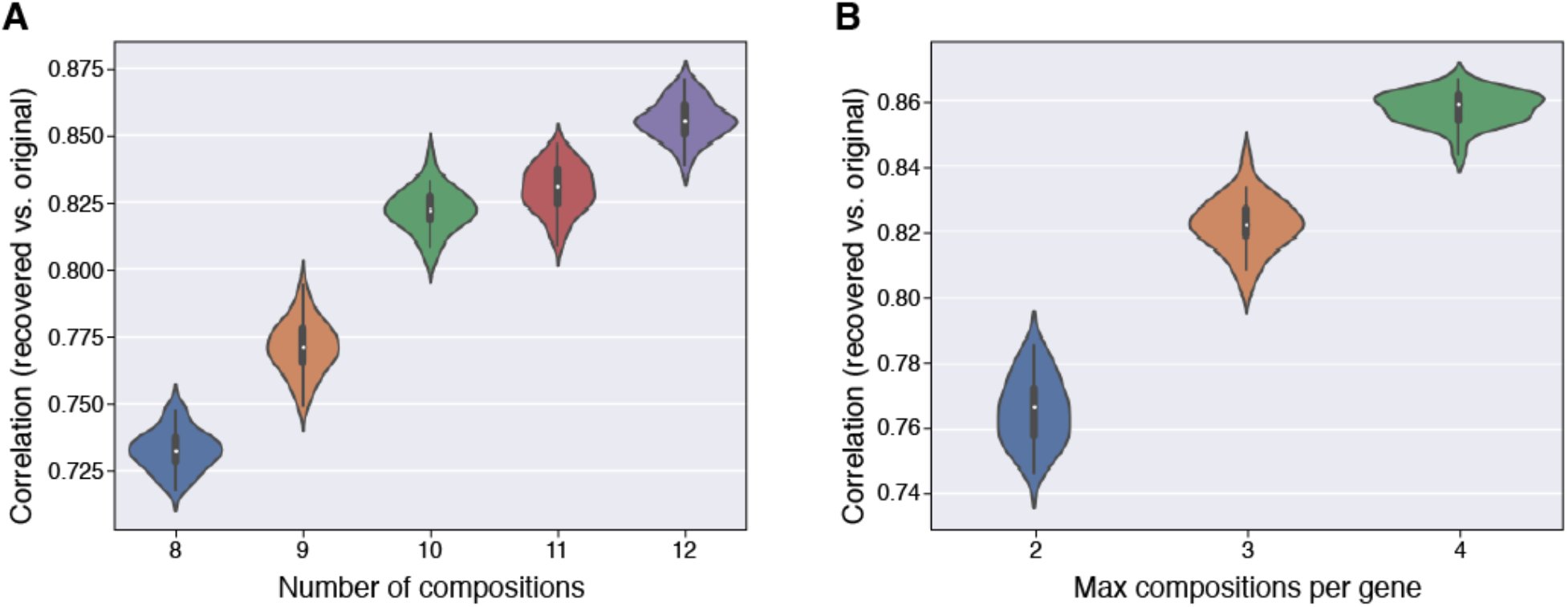
Evaluation of performance of simulated compositions. Distribution of Pearson correlation between the original and recovered expression levels of 37 genes in each cell (y axis) across simulation trials for different numbers of composite measurements (**a**), or for different measurement densities, set by the maximum number of measurements in which each gene was included (**b**). In (**a**) the maximum compositions per gene is 3, and in (**b**) the number of compositions is 10. Mini boxplots depict median (dots), inner quartiles (box), and 1.5x quartile range (whiskers).

**Supplementary Figure 4.**
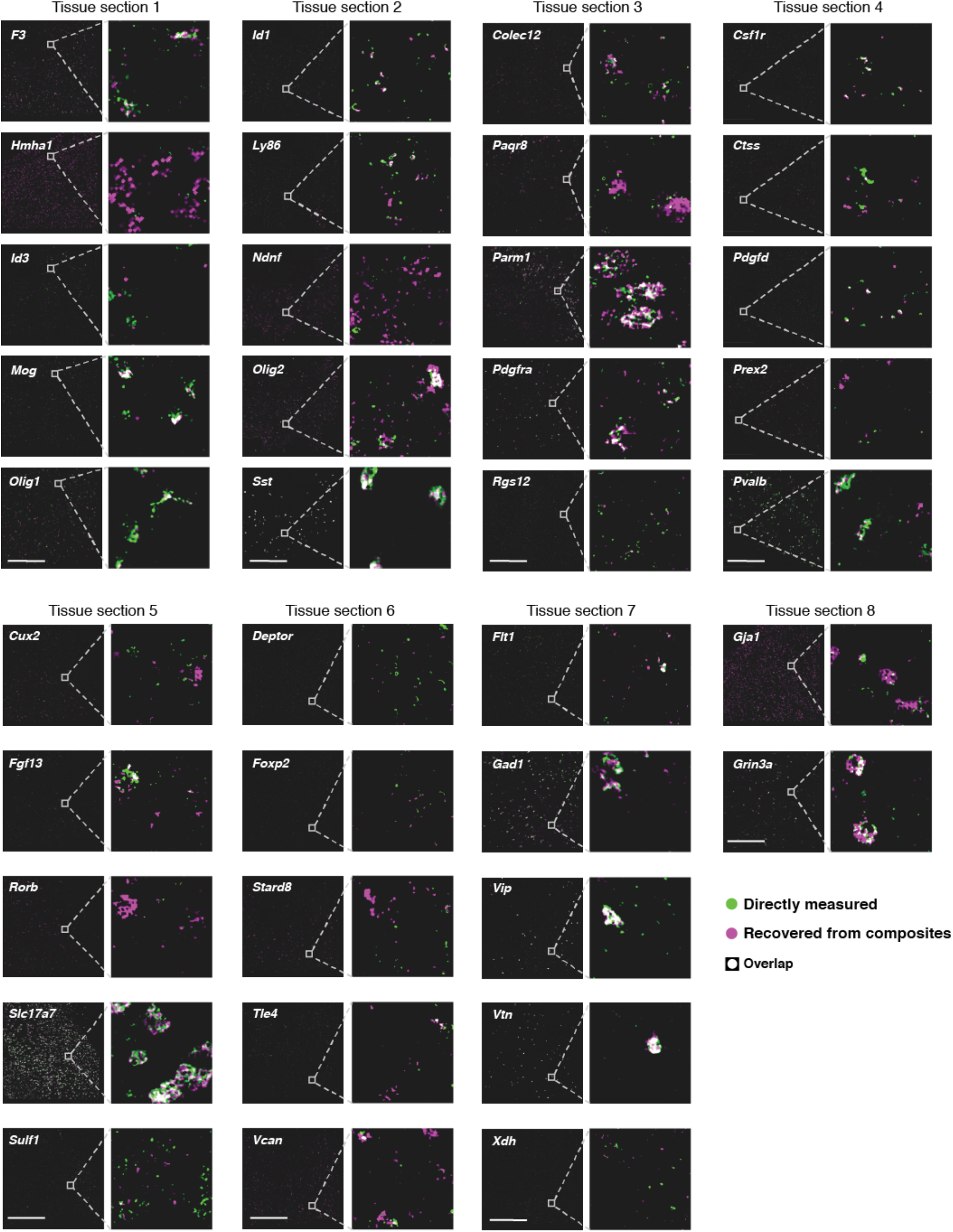
Autoencodeer based decompression successfully recovers accurate spatial patterns of individual genes compared to direct measurement on the same section. RNA images recovered by decompression with the segmentation free algorithm (magenta) and directly measured (green) in the same tissue section. White: images overlap exactly. Genes are grouped based on the section in which their direct measurements were made. Insets for all genes in a section show the same region, or an adjacent region if no cells for a given gene were present. Scale bar: 500um.

**Supplementary Figure 5.**
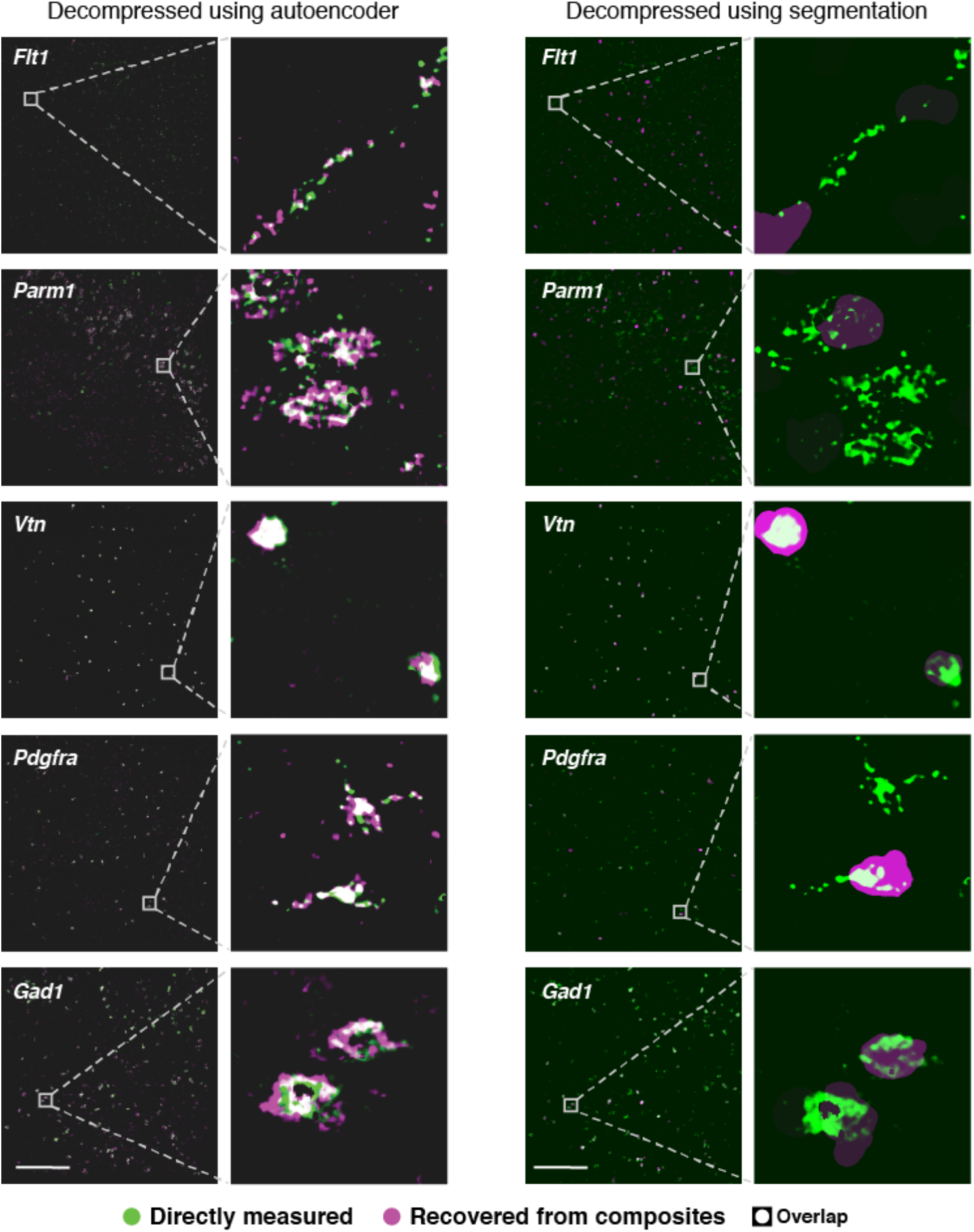
Comparison of autoencoding and segmentation-based decompression. Individual gene images recovered (magenta) using the autoencoding algorithm (left) or the segmentation based algorithm (right) are overlaid with direct measurement (green) of the genes in the same tissue sections (white: direct overlap). For segmentation-based decompression, the decompressed signal for each gene is projected uniformly over each segmentation mask. Scale bar: 500um.

**Supplementary Figure 6.**
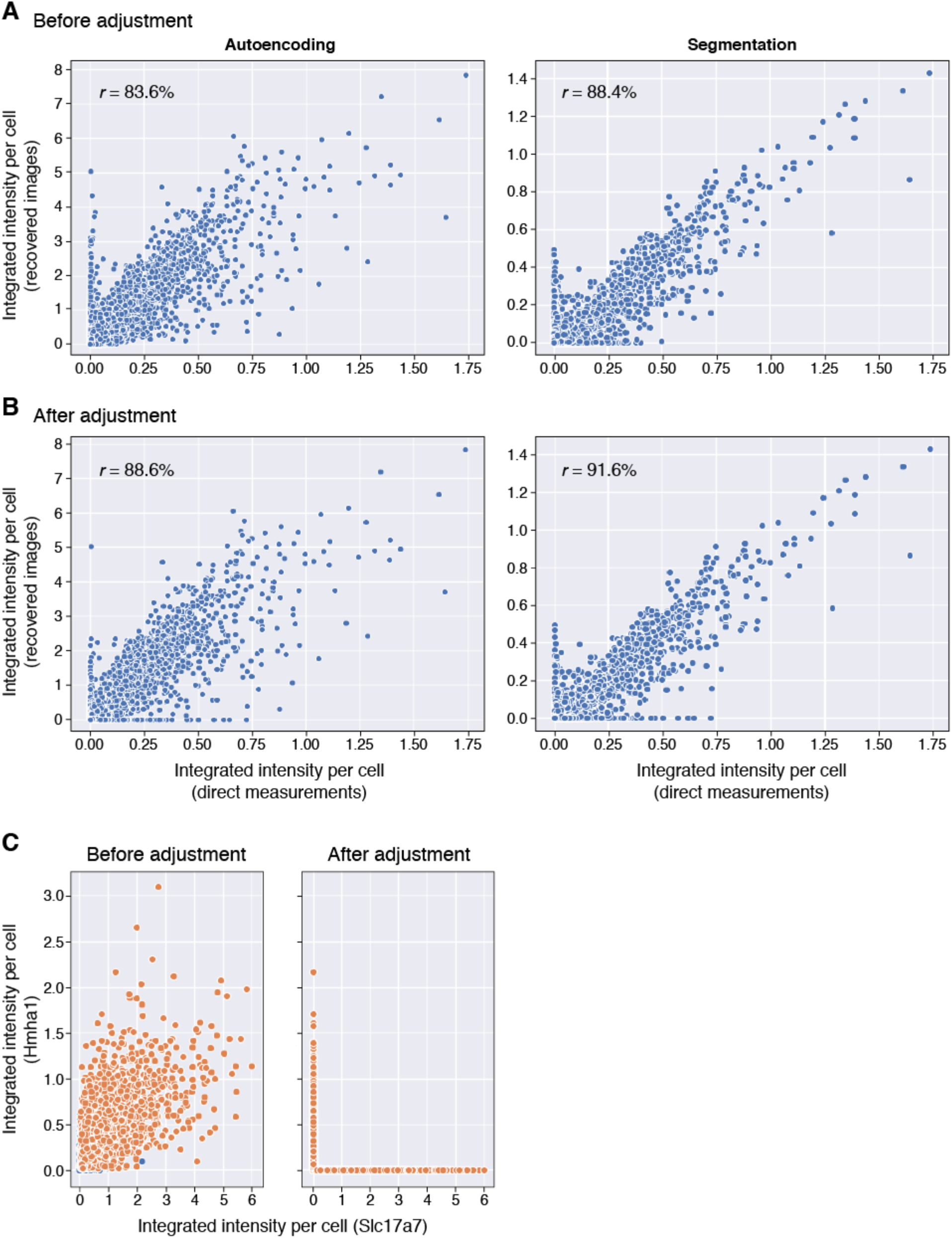
Evaluation of recovered signals before and after co-measurement adjustment. (**a,b**) Adjustment improves recovered signals. Integrated signal intensity for each gene in each cell (individual dots) from direct measurements (x axis) and from estimates recovered by the autoencoder decompressed images (y axis) either before (**a**) and after (**b**) co-measurement correction. (**c**) Example correction. Segmented cell intensities before (left) and after (right) correction for two co-measured genes (Hmha1 and Slc17a7) that were not correlated in snRNA-Seq.

**Supplementary Figure 7.**
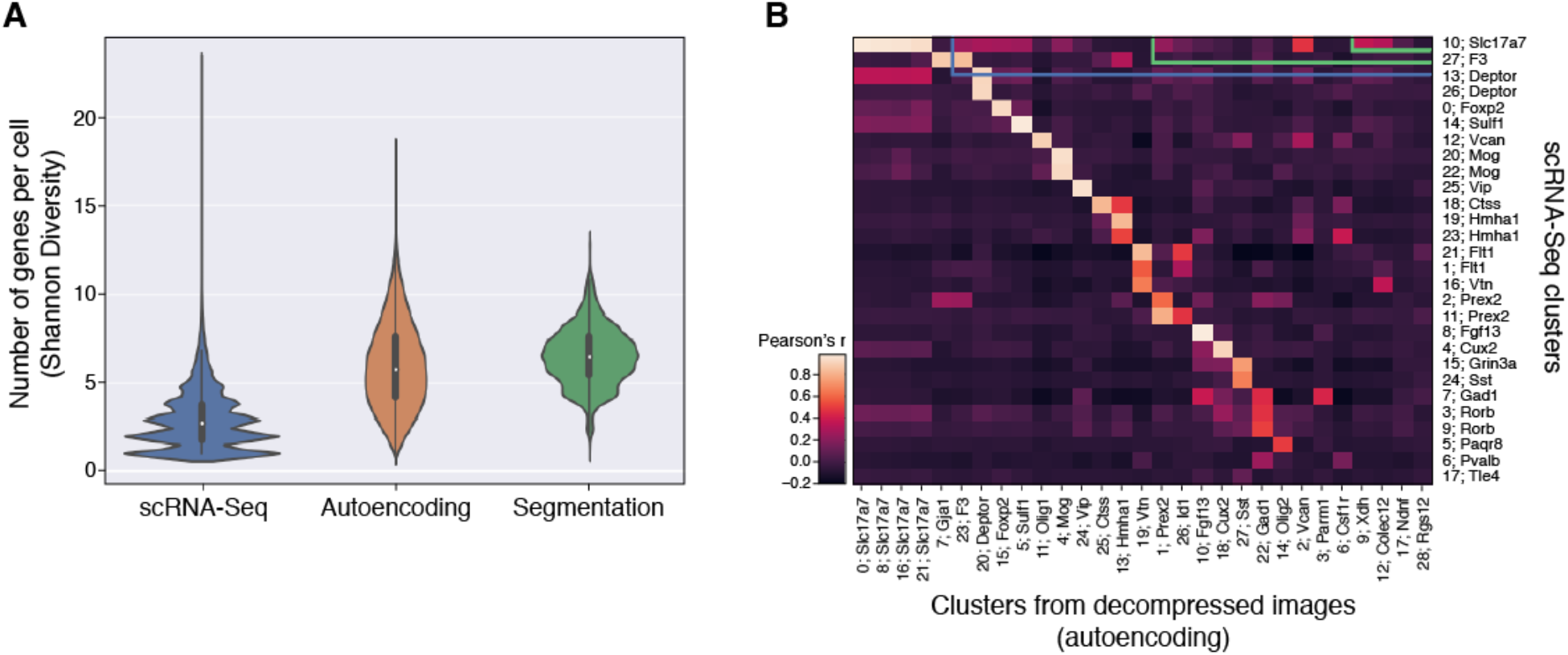
Evaluation based on genes per cell and cell clusters. (**a**) Distribution of expression diversity (effective number of genes expressed per cell out of 37 total; *y* axis) in snRNA-Seq, or based on recovered expression levels using autoencoding or segmentation-based decompression (*x* axis). Mini boxplots depict median (dots), inner quartiles (box), and 1.5x quartile range (whiskers). (**b**) Correspondence (Pearson’s correlation of mean gene expression; color bar) between cell clusters from snRNA-Seq (rows) and those found from *post hoc* segmentation of images recovered using the autoencoding algorithm (columns). One marker gene for each cluster is indicated.

**Supplementary Figure 8.**
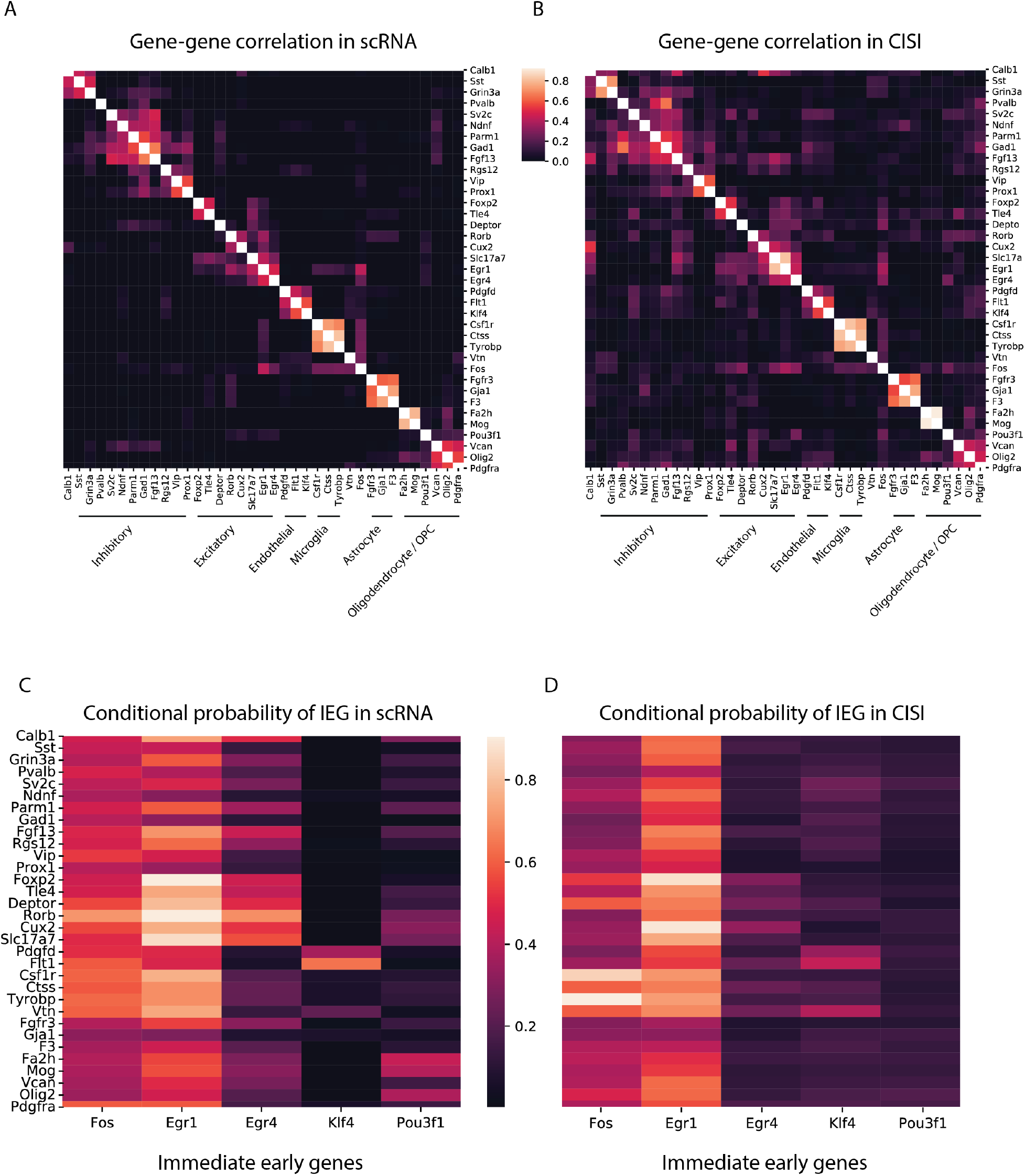
CISI recapitulates clusters and conditional probabilities from scRNA-Seq. (**a,b**) Consistent identification of cell type specific gene programs in scRNA-Seq and CISI. The correlation coefficient (colorbar) between pairs of genes (row and column labels) in scRNA-Seq (**a**) and decompressed CISI measurements (**b**). Rows and columns are clustered. Gene clusters of cell type specific markers are labeled by the respective cell type. (**c,d**) Consistent cell type expression patterns for IEGs in scRNA-Seq is CISI. Conditional probability (colorbars) of IEGs (columns) in cells that express a given gene (rows) in scRNA-Seq (**c**) and decompressed CISI (**d**) data.

